# Developmental interactions between oscillatory and spontaneous neural activity support improvements in inhibitory control from adolescence into adulthood

**DOI:** 10.64898/2026.11.05.764280

**Authors:** Abigail M. Beatty, Shane D. McKeon, Daniel J. Petrie, Finnegan J. Calabro, Ashley C. Parr, Will Foran, Beatriz Luna

**Affiliations:** Department of Bioengineering, University of Pittsburgh, PA, USA; Department of Human Development and Family Science, University of Georgia, Athens, GA, USA; Department of Psychiatry, University of Pittsburgh, Pittsburgh, PA, USA; Department of Psychology, University of Pittsburgh, Pittsburgh, PA, USA

**Author notes:** Correspondence: Abigail M. Beatty, Laboratory of Neurocognitive Development University of Pittsburgh, 121 Meyran Avenue, Pittsburgh, PA 15213.

## Abstract

Adolescence is a critical period of development marked by improvements in cognitive control driven by maturation of executive brain systems. Neural mechanisms supporting inhibitory control—the ability to suppress a prepotent response in favor of generating a goal-directed response—mature into adulthood in parallel with refinements in frequency-specific neural oscillations and broadband spontaneous activity, supporting more accurate and reliable performance on response inhibition tasks, such as the antisaccade oculomotor task. Prior work has established that the maturation of neural circuitry contributes to age-related decreases in spontaneous activity, supporting higher-fidelity transmission of task-relevant activity, such as frequency-specific neural oscillations, which are crucial for antisaccade performance. Using longitudinal electroencephalography (EEG) data from 146 participants (72 F, ages 10-34, 253 total sessions) collected while participants performed an antisaccade task, we sought to understand the interplay between frontal oscillatory and spontaneous activity during cognitive processing and how it supports developmental improvements in inhibitory control. We found that spontaneous activity decreased developmentally during the preparation of an inhibitory response, and that this was associated with reduced trial-to-trial variability in response latency, indicating that reductions in spontaneous activity may increase the reliability of inhibitory control processing. Furthermore, performance-related decreases in broadband spontaneous activity were associated with the suppression of preparatory high-beta oscillations in an age-dependent manner, such that this relationship became more pronounced through development. Together, these results illustrate how developmental enhancements in the coordination of frequency-specific oscillations and spontaneous activity during inhibitory response preparation support developmental improvements in inhibitory control, providing a potential mechanism through which inhibitory control becomes more accurate and reliable from adolescence into adulthood.

## 1. Introduction

Adolescence is a critical period of development known for protracted maturation of executive brain systems that underlie the stabilization and optimization of cognitive control into adulthood (Larsen and Luna, 2018; Tervo-Clemmens et al., 2023). One key aspect of cognitive control is inhibitory control—the ability to suppress a prepotent, reflexive response in favor of generating a goal-directed response (Aron, 2007; Luna et al., 2015). The neural circuitry underlying inhibitory control is available as early as infancy (Diamond and Goldman-Rakic, 1989; Johnson, 1995); however, the ability to consistently engage this circuitry emerges through the transition from adolescence to adulthood (Luna et al., 2015). Thus, the development of inhibitory control is not simply the emergence of the necessary neural circuitry, but the increasing ability to consistently engage these systems to support goal-directed behavior.

The antisaccade (AS) oculomotor task is widely used in both humans (Luna et al., 2008; Munoz et al., 1998) and non-human primates (Amador et al., 1998; Constantinidis and Luna, 2019) to probe the integrity of brain systems underlying inhibitory control and its development. Successful AS performance requires the suppression of a prepotent prosaccade (visually guided saccade) towards a suddenly appearing visual stimulus and the generation of a voluntary saccade to the mirror location of the visual stimulus (Munoz and Everling, 2004). Prosaccades reflect bottom-up, feedforward processing, which may bypass higher-order cognitive systems, whereas antisaccades require the engagement of brain systems related to top-down control to modulate the oculomotor system to adhere to task goals (Coe and Munoz, 2017; Cordones et al., 2013; Hwang et al., 2010; Ordaz et al., 2013). When preparing to inhibit a prosaccade towards the visual stimulus in the AS task, differences in activation and connectivity (compared to prosaccade) are seen in a distributed network of prefrontal, parietal, oculomotor, and subcortical regions involved in top-down control (Ettinger et al., 2008; Ford et al., 2005; Hwang et al., 2010). Engagement of top-down processes improves throughout adolescence and into adulthood, (Hwang et al., 2010; Ordaz et al., 2013) reflected in developmental decreases in response inhibition errors (Geringswald et al., 2020; Hwang et al., 2016; Ordaz et al., 2010; Parr et al., 2022), faster AS latencies (Hwang et al., 2016; Ordaz et al., 2010), and less variability in response latencies (Munoz et al., 1998).

Successful response inhibition relies on adequate preparation of an inhibitory response prior to the appearance of the visual cue (Everling et al., 1998; Everling and Munoz, 2000; Hwang et al., 2014; Ordaz et al., 2010; Schlag-Rey et al., 1997). Previous literature has shown that frequency-specific neural activity during the preparation of an antisaccade is predictive of successful AS performance in humans (Hwang et al., 2014) and non-human primates (Everling et al., 1998; Everling and Munoz, 2000; Schlag-Rey et al., 1997). Oscillations at different frequencies have been shown to support distinct phases of response inhibition. While alpha (∼8-12 Hz) activity from the frontal eye field (FEF) during antisaccade response preparation has been associated with correct responses in humans (Hwang et al., 2014) and non-human primates (Johnston et al., 2019), beta (∼13-34 Hz) activity in the dorsolateral prefrontal cortex (DLPFC) precedes FEF alpha activity, suggesting that beta oscillations exert top-down control over bottom-up signals (Hwang et al., 2016, 2014). Furthermore, the coupling between beta and alpha activity has been shown to strengthen developmentally (Hwang et al., 2016), providing a potential mechanism for developmental improvements in response inhibition via enhanced top-down control of preparatory signals. However, the mechanisms contributing to developmental enhancements in frequency-specific oscillations relevant for response inhibition remain incompletely understood.

One possibility is that in addition to frequency-specific oscillations, developmental changes in spontaneous activity—ongoing broadband neural activity that is not phase-locked to discrete task events— may alter the neural context in which task-related oscillations are generated. Spontaneous activity has been demonstrated to decrease developmentally in both resting-state (Miskovic et al., 2015; Ott et al., 2021) and an auditory steady-state task (McKeon et al., 2024a), which may increase the fidelity of frequency-specific oscillations involved in inhibitory response preparation, thus leading to more reliable behavioral performance. Importantly, ongoing activity can interact (e.g., during the preparatory period) with stimulus-locked neural responses (Wainio-Theberge et al., 2021), suggesting that spontaneous activity may shape the expression of task-related oscillatory dynamics. However, how spontaneous and frequency-specific activation patterns interact to support the maturation of cognitive control has not previously been investigated. Therefore, we sought to interrogate how age-related changes in spontaneous activity contribute to enhancements in frequency-specific oscillations, and how these two patterns of activity jointly shape the development of inhibitory control.

To address these questions, we leveraged longitudinal EEG data (253 total sessions) recorded during the preparatory period of the antisaccade task and contrasted this with the visually guided saccade (VGS) oculomotor task. We hypothesized that spontaneous activity during AS response preparation would decrease with age, allowing for higher-fidelity transmission of task-related oscillations supporting the age-related improvements in inhibitory control.

## 2. Methods

### 2.1. Participants

The full sample included data collected from 171 participants (87 assigned female at birth) between 10 and 34 years of age, with 66.5% identifying as White, 14.7% as Black, 7.6% as Asian, 0.6% as Hispanic, 8.8% as multiracial, and 1.8% not specifying. Participants were recruited as part of an accelerated longitudinal study design with up to 3 time points, approximately 18 months apart, for a total of 308 visits. Each time point consisted of 3 visits (a behavioral session, a 7T MRI scan, and an EEG session) each of which typically occurred within 1-2 weeks. Participants were recruited from the greater Pittsburgh area and were excluded if they had a history of loss of consciousness due to head injury, non-corrective vision problems, neurological disorders, personal or first-degree relative history of psychiatric disorders, or learning disabilities, or if they had characteristics that would disqualify them from MRI testing. All participants completed informed consent approved by the University of Pittsburgh IRB and were compensated for their involvement in the study. Aspects of MRI (Parr et al., 2025; Perica et al., 2022) and resting-state EEG (McKeon et al., 2024a, 2024b) have previously been reported from this line of inquiry.

### 2.2. Data acquisition and preprocessing

#### 2.2.1. EEG data

Electrooculogram (EOG) and high-impedance EEG were recorded concurrently with a Biosemi ActiveTwo 64-channel EEG system. Recording took place while stimuli were presented on a computer approximately 80 cm from the participant. The preprocessing pipeline was conducted using EEGLAB (Delorme and Makeig, 2004), with protocol that first down-sampled the data to 150 Hz from the original 512 Hz sampling rate. Data were referenced to external electrodes placed on the mastoids, given their distance from the scalp and low-signal recording of neural activity. The data were filtered with a non-causal finite impulse response (FIR) bandpass filter with high-pass cutoff frequency of 0.5 Hz and low-pass cutoff frequency of 70 Hz. Channel data were removed if they were identified as flatline (maximum tolerated flatline duration: 8 s) or noisy (defined as more than 5 standard deviations from the average channel amplitude), or if they contained short spontaneous bursts or incomplete segments of data. Removed channels were interpolated with data from surrounding electrodes. The data was re-referenced to the average reference for independent component analysis (ICA). Visual inspection of ICA components was performed to remove components that captured horizontal and vertical eye movements, eye blinks, and muscle artifacts. Finally, the resulting data was filtered with a non-causal FIR notch filter from 59 to 61 Hz to remove line noise. Continuous EEG data were segmented into epochs from 1 s before to 1 s after the onset of the preparatory fixation cross, providing a sufficient buffer to avoid edge artifacts during time-frequency analysis of the 500 ms preparatory window (Zhang, 2019). Epochs were rejected from analyses if their amplitude was more than 6 standard deviations from the single-channel mean amplitude or more than 2 standard deviations from the average channel mean amplitude. Protocol for preprocessing pipeline can be found on GitHub (https://github.com/LabNeuroCogDevel/EEG_preprocessing/tree/main).

#### 2.2.2. Oculomotor tasks

During EEG and EOG recordings, participants completed 40 trials each of the AS and VGS oculomotor tasks, administered in blocks with VGS trials presented first. The AS task is a well-validated measure used to probe the integrity of the circuitry underlying inhibitory control in both humans (Everling and Fischer, 1998; Hwang et al., 2016, 2014; Munoz et al., 1998; Ordaz et al., 2013; Parr et al., 2022) and non-human primates (Amador et al., 1998; Everling and Munoz, 2000; Schlag-Rey et al., 1997; Zhou and Constantinidis, 2017). The VGS task was administered as a control for potential developmental changes in core aspects of the oculomotor system beyond the engagement of the executive functions required for inhibitory control. Because the AS task has been described extensively elsewhere (Everling and Fischer, 1998; Luna et al., 2008), only a brief overview is provided here. Each AS trial began with a central fixation cross (gray central crosshair) for a variable period of 1-3 s, denoted as the intertrial interval (ITI). After the ITI, the central crosshair turned red, indicating the response preparation period during which participants should engage preparatory inhibitory processes. After 500 ms, the central crosshair was extinguished, and a visual stimulus (yellow dot) appeared ± 3 or 6 degrees of visual angle from the central crosshair. Participants were instructed to saccade to the mirror location of the visual stimulus as quickly and accurately as possible (Fig. 1A, left). During VGS trials, timing and structure were identical to AS trials; however, during the preparatory period the central crosshair turned green instead of red, cueing participants to prepare a saccade toward the upcoming visual stimulus rather than the mirror location (Fig. 1A, right).

**Fig. 1:**
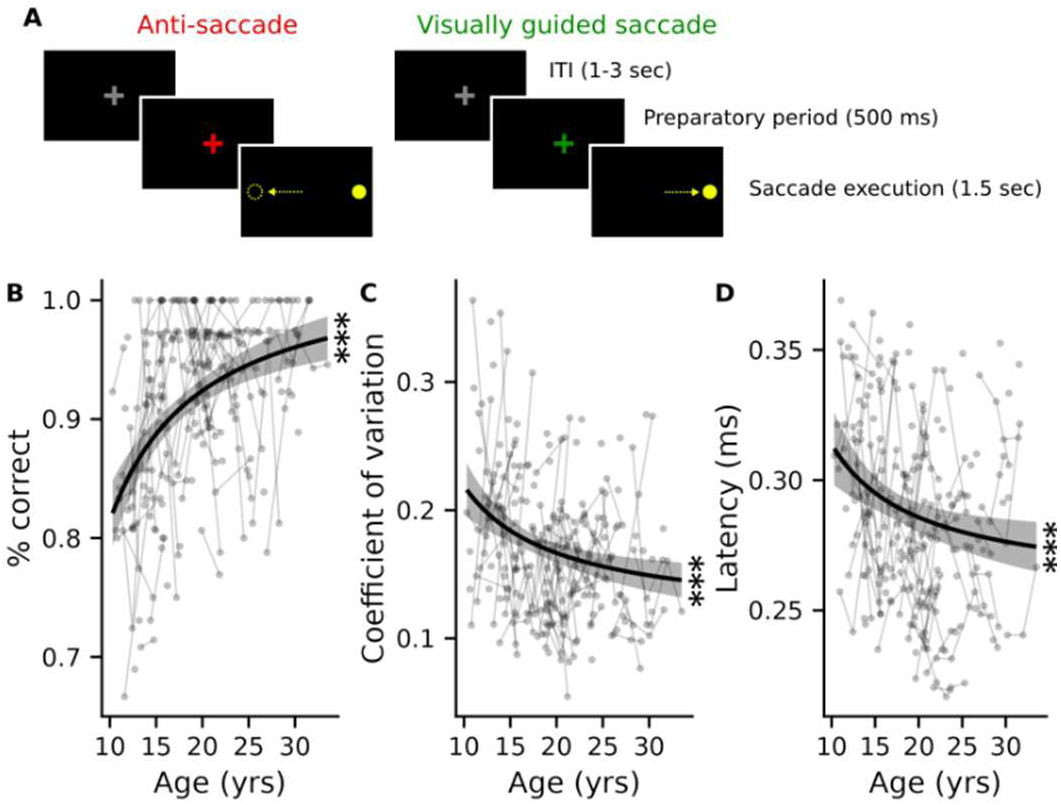
Antisaccade task and developmental performance. **A.** The antisaccade task to assess response inhibition **(left)** and the visually guided saccade task to control for potential developmental changes in oculomotor function **(right)**. The antisaccade task began with the participant maintaining fixation to the gray central cross hair. After a variable period of time (1-3 secs), the gray crosshair turned red, indicating to the participant to prepare an inhibition of a reflexive eye movement towards the stimulus. The central crosshair was extinguished after 500 ms, and the visual stimulus (yellow dot) appeared +/- 3 or 6 degrees from the central crosshair. The visually guided saccade had the same timing of events; however, the central crosshair turned green, indicating an eye movement towards the visual stimulus was required. Developmental changes in **B.** AS performance (correct response rate), **C.** trial-to-trial latency variability for correct trials, and **D.** mean response latency on correct trials. Gray circles represent raw empirical measures of performance across age, where the same subject’s data points are connected with thin gray lines. Solid black lines indicate the linear regression predicted trend of behavior across age, controlling for sex, with the light gray region showing the standard error. (*** = p < 0.001).

### 2.3. Data analysis

#### 2.3.1 Behavioral analyses

Horizontal EOG channels were used to track eye position during the tasks. A calibration routine was performed prior to the start of the tasks to calibrate the participant’s eye position with the EOG signal. Automated eye calibration and scoring were performed using custom software available on GitHub (https://github.com/LabNeuroCogDevel/7TBrainMech_scripts/tree/master/eeg/eog_cal). Correct AS trials were defined as those in which the first eye movement during the response period was generated toward the mirror location of the stimulus. Incorrect AS trials were defined as trials in which the saccade remained directed toward the stimulus, and error-corrected trials were defined as those in which the initial saccade toward the visual stimulus was followed by a corrective saccade to the mirror location. Trials in which no eye movement was generated were defined as non-response trials and were excluded from analysis. AS trials with latencies less than 100 ms (express saccade latency), which represent reflexive, anticipatory responses, were excluded from analysis. To ensure analyses only included participants that understood the task, we excluded individuals with fewer than 20 AS trials scored as either correct or error-corrected (Parr et al., 2022).

After preprocessing and application of exclusion criteria, analyses included 253 EEG sessions from 146 participants (72 assigned female at birth) between 10.21 and 33.52 years old. A two-sided Kolmogorov-Smirnov test did not detect significant differences in the age distribution after exclusion (*D* = 0.04, *p* = 0.95), reflecting that the exclusion criteria did not disproportionately exclude younger ages (Fig. S1A).

Response latency for each trial was defined as the time difference between the stimulus onset and initiation of an eye movement (when velocity was greater than or equal to 30 degrees/s), and average latency on correct trials was calculated per participant. The coefficient of variation, which reflected latency variability, was defined as the standard deviation of response latencies on correct trials divided by the average latency on correct trials. Finally, correct response rate was defined as the number of correct trials divided by the total number of correct and error corrected trials (Parr et al., 2022).

#### 2.3.2. EEG analyses

During the AS task, preparatory processes prior to the appearance of the visual stimulus have been shown to predict correct inhibition of the reflexive response in both non-human primates (Johnston et al., 2019) and humans (Hwang et al., 2014). The overall goal of this study was to understand developmental changes in the mechanisms of correctly engaging inhibitory control processes. Therefore, we performed all analyses during the preparatory period of the VGS and AS tasks and included only correct AS trials. The code used in analyses can be found on GitHub (https://github.com/AMB895/AS-EEG).

##### 2.3.2.1. Time-frequency reconstruction

EEG data time-locked to the onset of the fixation cross for both oculomotor tasks were extracted from each channel, 1 s before to 1 s after fixation cue. For each trial, EEG data *s* at time *t* were convolved with a complex Morlet wavelet *G* centered at frequency *f*, resulting in a complex time-frequency reconstruction *w* of the EEG data (Hwang et al., 2016).

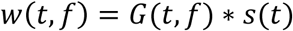

Where

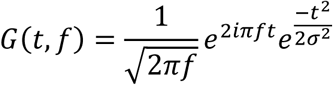

Frequency *f* spanned the 3-70 Hz range in linear steps with a 2.2 Hz increment. The Gaussian width *σ* was defined by the number of cycles *n*, which increased linearly with frequency *f* from 3 to 35 cycles. The full width at half maximum (FWHM) ranged from 187 to 373 ms, decreasing with wavelet frequency *f* (Cohen, 2019).

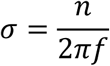

Power *P* was computed by squaring the absolute value of the complex time-frequency EEG data w, for each channel and trial *n*.

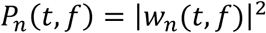

The event-related spectrum (ERSpec) across *N* trials is defined as (Grandchamp and Delorme, 2011):

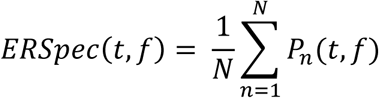

Event-related spectral perturbation (ERSP) is a measure of the change in spectral power from baseline over time. Baseline power *μ* was defined as the period from t_1_=700 to t_2_=400 ms prior to the onset of the preparatory period, averaged across *N* trials and *M* time points, consistent with previous studies (Grandchamp and Delorme, 2011; Hwang et al., 2016).

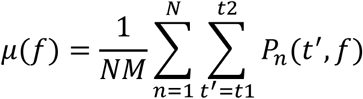

ERSP was computed by normalizing the mean ERSpec across trials by the baseline power spectrum and converting to decibels (Grandchamp and Delorme, 2011).

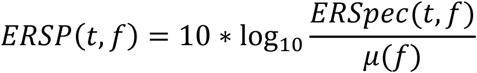

Finally, the ERSP data was averaged across frontal electrodes (F1, F2, F3, F4, F5, F6, F7, F8, Fz), which corresponded to frontal brain regions with a well-established role in inhibitory control (Bells et al., 2020; Brown et al., 2007; Hwang et al., 2010; Luna et al., 2015; Ordaz et al., 2013) for correct AS and VGS trials. To determine whether our findings were specific to frontal processes, we ran separate analyses on parietal electrodes (P1, P2, P3, P4, P5, P6, P7, P8, P9, P10, Pz). The event-related spectral changes from baseline can be characterized as an event-related desynchronization (ERD; decrease in spectral power) or an event-related synchronization (ERS; increase in spectral power) and are time-locked, but not phase-locked, to the onset of the preparatory period (Zhang, 2019).

##### 2.3.2.2. Task-related evoked and spontaneous activity

Neural activity can be characterized as evoked (stimulus-locked) or spontaneous (non-stimulus-locked or trial-to-trial fluctuations). Studies investigating evoked and spontaneous activity during tasks generally consider pre-stimulus activity as spontaneous and post-stimulus activity as evoked. However, this approach may miss how ongoing spontaneous activity impacts stimulus-evoked activity (Wainio-Theberge et al., 2021). To probe ongoing spontaneous activity during presentation of the preparatory cue, we used an established method (McKeon et al., 2024a) to calculate evoked and spontaneous power during the preparatory period of the AS task. Power spectra were computed using the method described in 2.3.2.1 and averaged across time. Evoked power was computed by averaging EEG epochs in the time domain across trials (the event-related potential) and then reconstructing this event-related potential in the frequency domain, which isolates activity that is time- and phase-locked to the onset of the preparatory period. Total power was computed by reconstructing each trial’s data in the frequency domain, maintaining all activity during the preparatory period regardless of whether it was consistently time- and/or phase-locked to the visual cue across trials. Spontaneous power was calculated by subtracting the evoked power spectrum from each trial’s total spectrum and averaging across trials, providing a measure of the trial-to-trial fluctuations in neural activity, this is, how much of the power present across trials varies across trails relative to the task timing. The evoked power spectrum characterizes activity that is consistent across trials in the time domain, thus trial-to-trial fluctuations in activity can be separated from overall activity by subtracting the evoked power spectrum from each trial’s total power spectrum. Finally, broadband (3-70 Hz) spontaneous and evoked power were computed by averaging the power spectrums across frequencies and frontal electrodes (F1, F2, F3, F4, F5, F6, F7, F8, Fz). As with ERSP analyses, we ran separate analyses on parietal electrodes (P1, P2, P3, P4, P5, P6, P7, P8, P9, P10, Pz) to determine if our findings were specific to frontal processes. Visualization of this method can be found in McKeon et al, 2024a.

#### 2.3.3. Statistical analyses

All statistical models were conducted in R (version 4.5.1; R Core Team, 2025). We accounted for the longitudinal structure of the data by using linear mixed-effects models with participant-level random intercepts for all analyses. To account for nonlinear developmental effects, separate models tested for effects of inverse age, which reflects rapid change during adolescence that decelerates into adulthood (Luna et al., 2004), and linear age. To confirm which model best described developmental effects, we performed an AIC analysis comparing the model fits with inverse age against linear age for age-trajectories of AS behavioral performance (Supplemental Table 1). Inverse age provided a better fit for the developmental trajectory of all three AS behavioral outcomes; therefore, we used inverse age in subsequent analyses. Outlier detection was performed separately for each analysis by fitting a linear model with age and relevant predictors for that analysis (e.g., ERSP, spontaneous/evoked power) and excluding observations with residuals exceeding ±2 standard deviations from the mean. Following outlier removal, all primary analyses were fit using linear mixed-effects models with participant-level random intercepts and sex was included as a covariate. Across analyses, false discovery rate (FDR) correction was applied to families of related tests to account for multiple comparisons, unless otherwise noted.

We first examined age-related changes in AS behavioral performance using measures of correct response rate, mean response latency, and trial-to-trial latency variability. We tested for age-by-sex interactions, which are reported in the Supplemental Material. Developmental trajectories of frontal broadband spontaneous and evoked power were characterized separately during both oculomotor tasks. Effects of (inverse) age, task condition, and their interaction were evaluated to determine whether EEG measures were significantly modulated by cognitive demands during the AS task. We next examined associations between spontaneous and evoked power and measures of AS performance (correct response rate, average latency, and latency variability). These tests were conducted separately for the AS and VGS task conditions. To determine whether these associations were specific to frontal brain regions, we compared broadband spontaneous and evoked power between frontal and parietal electrodes (reported in the Supplemental Material) and examined associations between behavioral performance and parietal broadband spontaneous and evoked power to assess the regional specificity of observed brain– behavior relationships. Finally, using ERSP clusters that were significantly related to AS behavior (method outlined in section below), we assessed the association between oscillatory activity and broadband spontaneous and evoked power by testing for significant main effects of average cluster ERSP, as well as significant average cluster ERSP-by-(inverse) age interactions on broadband spontaneous and evoked power.

##### 2.3.3.1. Characterizing ERSP cluster specific to age-related changes in cognitive control aspects of the AS task and their association with AS performance

To characterize developmental and task-related differences in frontal ERSP activity, linear mixed-effects models were fit at each time-frequency point. Fixed effects included (inverse) age, task condition (AS vs. VGS), and their interaction. These models were estimated using the fitlme() function in MATLAB.

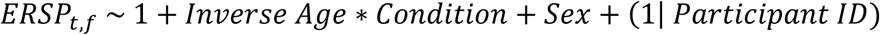

Significant ERSP clusters were identified using a threshold-free cluster enhancement (TFCE) algorithm implemented by an EEGLAB plug-in, LIMO Tools, which, unlike traditional clustering methods, does not require a predefined cluster threshold (Maris and Oostenveld, 2007; Smith and Nichols, 2009). The TFCE algorithm takes the t-values from the linear model and enhances each value based on strength and size of surrounding data. Significant clusters were determined by permuting the t-value maps 1000 times, generating a null distribution of maximum TFCE values and setting the enhanced t-value threshold at p < 0.01. If a cluster had an area fewer than 70 time-frequency points, it was removed from analyses (Mensen and Khatami, 2013). A visualization of the TFCE cluster method is shown in Fig. S2. We were interested in developmental changes in activity specific to engaging response inhibition processes, indexed by significant ERSP clusters from the (inverse) age-by-condition interaction. The average ERSP from significant (inverse) age-by-condition time-frequency clusters was extracted for each participant for use in subsequent analyses.

Using a linear mixed effects model, we assessed the associations between average ERSP from *S* significant clusters and AS behavior, controlling for sex and (inverse) age.

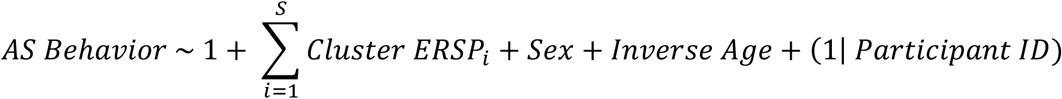

To determine whether these clusters were specific to frontal brain regions, we performed the same analyses on average parietal ERSP from P-row electrodes. Only clusters significantly associated with AS behavior were included in subsequent analyses.

## 3. Results

### 3.1. Developmental improvements in antisaccade performance

Linear mixed-effects models revealed that AS behavioral performance improved with age, indicated by an increased correct response rate (β = -2.19, *t* = -7.04, *p* < 0.001; Fig. 1B), reduced trial-to-trial latency variability, (β = 1.05, *t* = 4.47, *p* < 0.001; Fig. 1C) and faster latencies on correct trials (β = 0.56, *t* = 3.55, *p* < 0.001; Fig. 1D). These findings are consistent with prior work in both humans (Hwang et al., 2016; Ordaz et al., 2013; Parr et al., 2022) and non-human primates (Constantinidis and Luna, 2019). There were no significant sex differences in correct response rate (β = 0.006, *t* = 0.60, *p* = 0.56), latency variability (β = 0.007, *t* = 1.01, *p* = 0.56) or average latency (β = 0.003, *t* = 0.59, *p* = 0.56). Age-by-sex interactions on AS behavioral performance are reported in the Supplemental Material (Fig. S3).

### 3.2. Developmental improvements in antisaccade behavioral performance related to age-related decreases in broadband spontaneous and evoked power

Linear mixed-effects models revealed developmental decreases in frontal broadband spontaneous power (β = 108.20, t = 14.54, p < 0.001; Fig. 2A, top), with significant main effects of condition (β = -0.95, t = -3.04, p = 0.005; Fig. 2A, top) and sex (β = -0.72, t = -2.72, p = 0.007). Frontal spontaneous power also exhibited a significant age-by-condition interaction (β = 18.37, t = 3.34, p = 0.002; Fig. 2A, top) in which the developmental trajectory of VGS power was steeper than that of AS power, reflecting age-dependent modulation of spontaneous activity during inhibitory control trials.

**Fig. 2:**
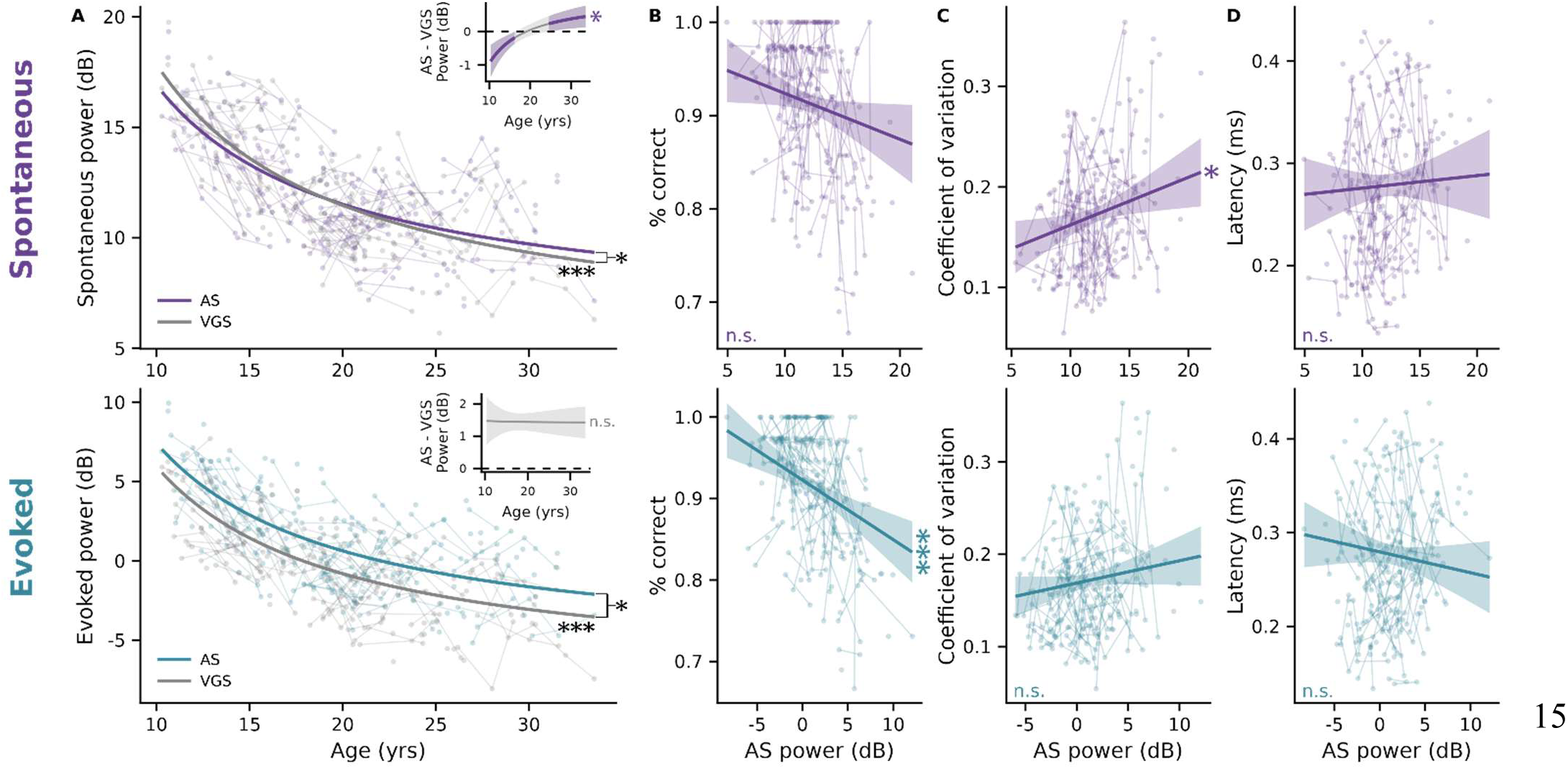
Developmental changes in spontaneous and evoked power and associations with AS behavior. **A (top).** Age-related trajectories of broadband spontaneous power for AS (purple) and VGS (gray) trials. The inset shows the developmental trajectory of the difference between AS and VGS spontaneous power, illustrating the age-dependent modulation of spontaneous power, such that children suppress spontaneous power on AS trials more than adults (relative to spontaneous power on VGS trials. **A (bottom).** Age-related trajectories of broadband evoked power for AS (blue) and VGS (gray) trials. The inset shows the developmental trajectory of the difference between AS and VGS evoked power, where there is no age-dependent modulation. Circles represent raw empirical measures of power across age, where the same subject’s data points are connected with thin lines. Solid lines indicate the linear regression predicted trend of power across age, controlling for sex. Associations between spontaneous **(top)** and evoked **(bottom)** power on AS trials and **B.** correct reponse rate, **C.** trial-to-trial latency variability, and **D.** mean reponse latency. Circles represent raw empirical measures of behavior across AS power (spontaneous = purple; evoked = blue), where the same subject’s data points are connected with thin lines. Solid lines indicate the linear regression predicted trend of behavior across power, controlling for age and sex. (n.s. = not significant; * = p < 0.05; ** = p < 0.01; *** = p < 0.001).

Frontal broadband evoked power also decreased significantly with age (β = 138.86, t = 14.05, p < 0.001; Fig. 2A, bottom), with significant main effects of condition (β = -1.36, t = -2.81, p = 0.005; Fig. 2A, bottom) and sex (β = -0.93, t = -2.95, p = 0.007). These effects were consistent across AS and VGS conditions (no age-by-condition interaction; β = -1.84, t = -0.22, p = 0.83; Fig. 2A, bottom). Age-by-sex interactions on broadband spontaneous and evoked power are reported in the Supplemental Material (Supplemental Table 2).

Age-related decreases in frontal spontaneous and evoked power showed distinct associations with behavior. Decreased frontal evoked AS power was associated with improved AS correct response rate (β = -0.007, t = -4.31, p = 0.002; Fig. 2B, bottom), whereas decreased frontal spontaneous AS and VGS power were associated with reduced variability (AS: β = 0.005, t = 2.55, p = 0.05; Fig. 2C, top. VGS: β = 0.005, t = 2.53, p = 0.05; Fig. S4B, top). Neither spontaneous nor evoked AS or VGS power was associated with mean AS latency (AS: Fig. 2D; VGS: Fig. S4C, top).

To determine if associations between broadband spontaneous or evoked power and AS performance were specific to frontal processes, we first assessed differences in frontal and parietal broadband spontaneous and evoked power. There was greater frontal spontaneous power compared to parietal spontaneous power during AS trials (β = -1.34, t = -3.83, p = 0.006), with a significant interaction between age and row (β = 15.60, t = 2.55, p = 0.04), in which parietal spontaneous power decreased more steeply with age. There was no main effect of row on evoked power during AS trials (β = -0.61, t = -1.31, p = 0.38); however, there was an interaction effect (β = 18.40, t = 2.25, p = 0.05). There were no significant differences in VGS spontaneous or evoked power between frontal and parietal electrodes. Additional interactions tested are reported in the Supplemental Material. Across measures of parietal AS and VGS power, only evoked AS power showed associations with behavior with reductions in power associated with improved correct response rate (β = -0.006, t = -3.75, p < 0.001; Fig. S4A, bottom).

### 3.3. ERSP clusters reflecting age-related changes in inhibitory control are related to antisaccade behavioral performance

Time-frequency analyses identified five frontal (Fig. 3A) and six parietal (Fig. S5A) ERSP clusters during the preparatory period that showed an age-by-condition interaction, reflecting age-related change in the cognitive control aspects of the AS task (Table 1). Of the frontal clusters, average AS ERSP from the mid-gamma (44-50 Hz) and high-beta (30-32 Hz) clusters showed significant associations with AS performance. ERSP values less than zero indicate suppression (“desynchronization”) of activity from baseline, whereas values greater than zero indicate activation (“synchronization”) from baseline. Reduced suppression of AS mid-gamma was associated with an improved correct response rate (β = 0.03, t = 2.99, p = 0.003; Fig. 3D, top) and lower trial-to-trial latency variability (β = -0.02, t = -3.11, p = 0.002; Fig. 3E, top), whereas increased suppression of AS high-beta was specifically associated with reduced latency variability (β = 0.02, t = 2.90, p = 0.004; Fig. 3E, bottom).

**Fig. 3:**
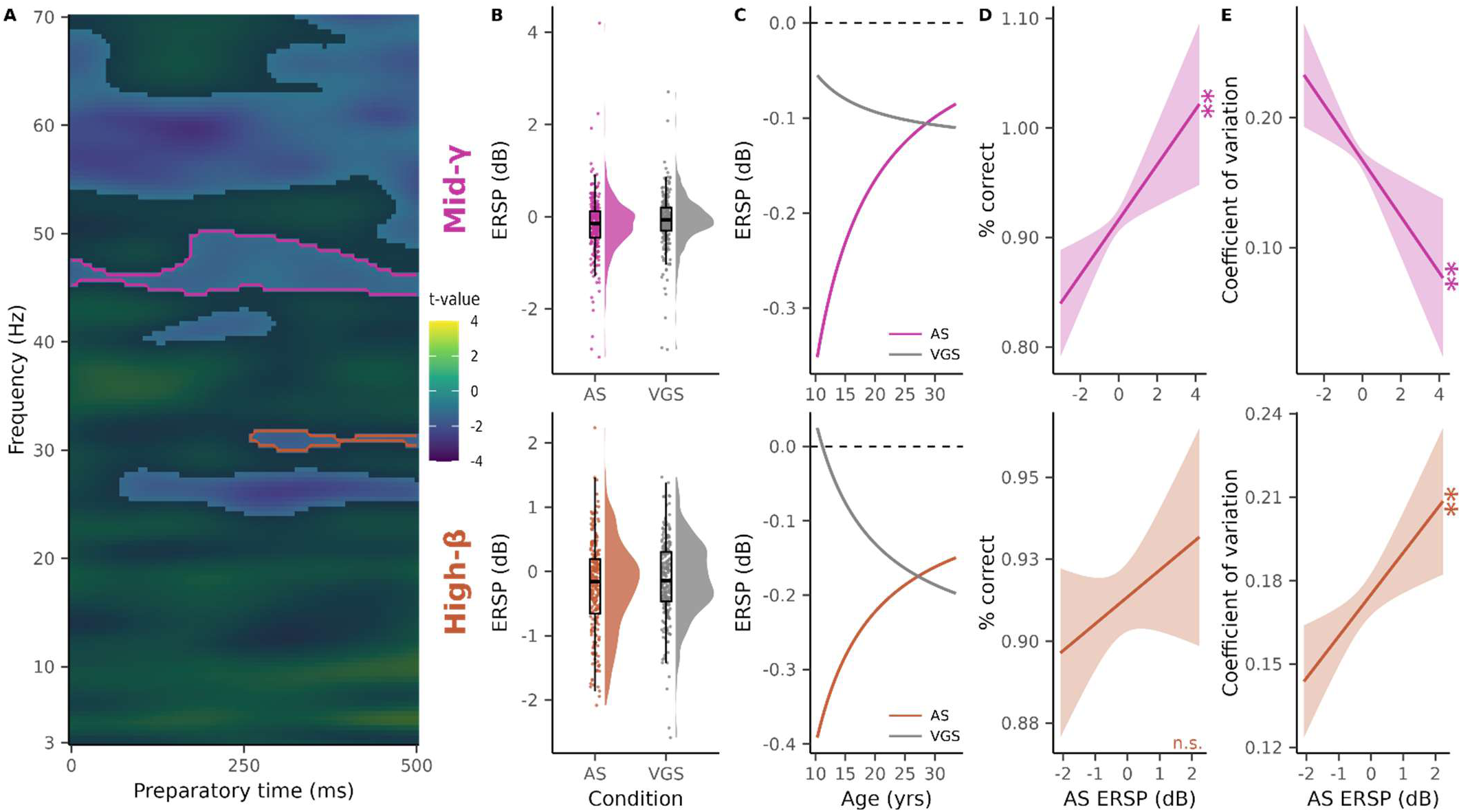
Frequency-specific oscillations associations with AS performance and reliability. **A.** Significant frontal ERSP clusters from the interaction (age * condition) term in the linear mixed effects model from section 2.3.3.3, where the x-axis is time throughout the preparatory period in milliseconds, the y-axis is frequency and the color bar (z-axis) is the t-value. Five significant clusters where identified; however, only mean ERSP from the mid-gamma cluster (pink) and high-beta cluster (orange) were related to AS behavior. The distribution of mean ERSP from the **B. (top)** mid-gamma cluster (AS: pink; VGS: gray) and **B. (bottom)** high-beta cluster (AS: orange;VGS: gray). Due to the large variability in the clusters’ mean ERSP, linear regression model predictions are shown in subsequent plots (C-E), controlling for sex (C-E) and age (D-E). Age-related trajectories of mean **C. (top)** mid-gamma AS (pink) and VGS (gray) ERSP and **C. (bottom)** high-beta AS (orange) and VGS (gray) ERSP, where the black dashed line represents no change in power from baseline, thus negative ERSP indicates a suppression in power from baseline and a positive ERSP indicates an activation in power from baseline. Associations between correct response rate and **D. (top)** mid-gamma and **D. (bottom)** high-beta AS ERSP. Associations between latency variability and **E. (top)** mid-gamma and **E. (bottom)** high-beta AS ERSP. (n.s. = not significant; * = *p* < 0.05; ** = *p* < 0.01).

**Table 1:** Frequency-specific oscillations associations with AS performance and reliability. Summary of AS ERSP clusters and their associations with behavior. F-row corresponds to electrodes associated with frontal brain regions.

| Electrodes | Cluster |  |  | Correct Response Rate |  | Coefficient of variation |  |
|---|---|---|---|---|---|---|---|
|  | Frequency (Hz) |  | Time (ms) | Estimate (β) | p | Estimate (β) | p |
| F-row | Mid-gamma | 44.4-50.2 | 0-500 | 0.03 | 0.003 | -0.02 | 0.002 |
|  | High-gamma | 48.9-70 | 0-500 | -0.006 | 0.49 | 0.0008 | 0.9 |
|  | Mid-beta | 24.1-28.2 | 73.3-500 | -0.005 | 0.59 | 0.008 | 0.23 |
|  | Low-gamma | 39.9-43 | 106.7-293.3 | -0.003 | 0.72 | 0.005 | 0.39 |
|  | High-beta | 30-31.8 | 260-500 | 0.008 | 0.21 | 0.02 | 0.004 |

### 3.4. High-beta antisaccade ERSP related to broadband spontaneous and evoked antisaccade power at different developmental stages

We next tested whether the two AS ERSP clusters that were significantly related to AS behavior (high-beta and mid-gamma) were associated with broadband AS power (spontaneous and evoked), and how these relationships varied by age. High-beta ERSP displayed a significant interaction with age on broadband spontaneous power (β = -19.58, t = -2.69, p = 0.03; Fig. 4B, top) and evoked power (β = -23.26, t = -2.30, p = 0.04; Fig. 4B, bottom). Johnson-Neyman plots were used to visualize these interactions, which demonstrated that reduced high-beta ERSP was related to lower spontaneous power at ages above 17.4 years old (Fig. 4C, top), whereas reduced high-beta ERSP was related to higher evoked power at ages below 15.5 years old (Fig. 4C, bottom). This pattern may suggest a shift in the mechanisms employed to correctly complete the AS task between adolescence and adulthood. Mid-gamma ERSP displayed no significant interactions with age on broadband spontaneous (β = -1.22, t = -0.15, p = 0.88) or evoked (β = -18.92, t = -1.61, p = 0.14) power.

**Fig. 4:**
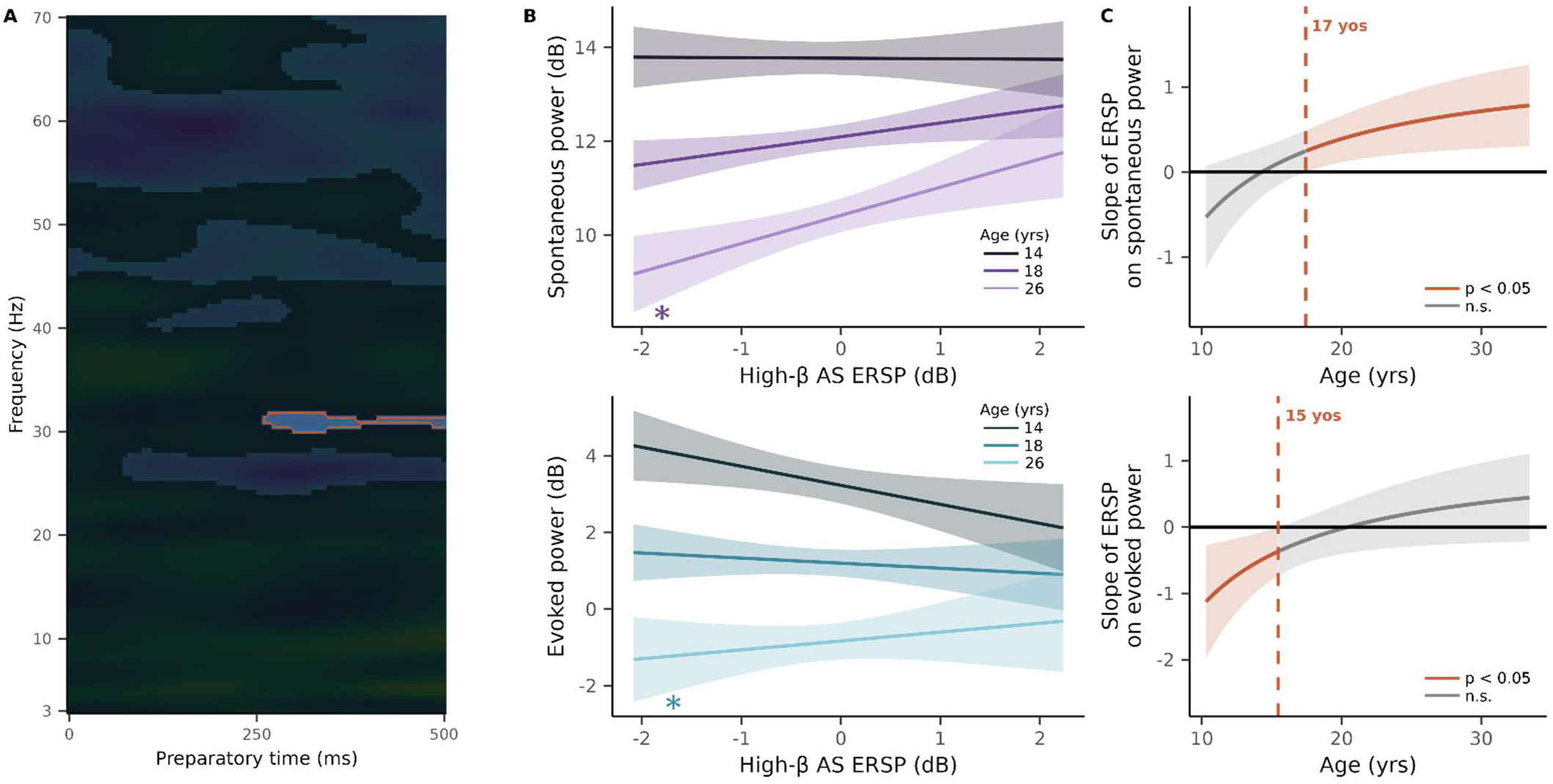
Age-dependent modulation of mid-beta oscillations on spontaneous and evoked power. **A.** High-beta ERSP cluster. Age-dependent modulation of high-beta AS ERSP on **B. (top)** spontaneous (purple) and **B. (bottom)** evoked (blue) AS power at representative ages (-1 standard deviation, mean, and +1 standard deviation): 14 years-old (dark purple/ blue), 18 years-old (medium purple/ blue), and 26 years-old (light purple/ blue). Johnson-Neyman plots showing the ages when the modulation of high-beta AS ERSP on **C. (top)** spontaneous and **C. (bot\tom)** evoked AS power is significant. (n.s. = not significant; * = *p* < 0.05)

## 4. Discussion

In this study, we used longitudinal EEG data during the performance of an oculomotor response inhibition task (the antisaccade task) to characterize how age-related changes in frequency-specific oscillatory activity contribute to overall shifts in broadband spontaneous and evoked activity, which may facilitate developmental improvements in inhibitory control. Consistent with prior findings, we found that AS performance improved developmentally, becoming more reliable with faster response latencies. We found that age-related reductions in frontal spontaneous activity during the preparatory period allowed for increases in AS reliability, whereas developmental decreases in frontal evoked power were linked to improved AS performance. Furthermore, we found that frontal oscillatory activity related to the developmental changes in inhibitory response preparation in the high-beta (30-32 Hz) and mid-gamma (44-50 Hz) ranges was associated with AS behavior: reductions in mid-gamma ERD supported improved AS performance and reliability and increases in high-beta ERD supported improved AS reliability. Of particular interest, frontal high-beta oscillations exhibited age-dependent modulation of preparatory spontaneous and evoked power. In adults, greater high-beta ERD was related to reductions in spontaneous power whereas, in adolescents greater high-beta ERD was associated with increased evoked power. Together, these results suggest joint contributions of broadband spontaneous and evoked power and frequency-specific oscillations on inhibitory control performance, which may vary with age to support the developmental refinements in inhibitory control.

We show that developmental decreases in spontaneous activity during both the AS and VGS task correlated with reduced AS latency variability. Spontaneous, ongoing activity represents a baseline of neural activity (Northoff et al., 2023). While spontaneous activity is commonly studied during rest (Uddin, 2020), recent evidence suggests that spontaneous activity persists during a task and can affect a stimulus-evoked response (Liu et al., 2022; Northoff et al., 2023; Wainio-Theberge et al., 2021). Adolescence is thought to represent a critical period of plasticity (Larsen and Luna, 2018), in which circuits undergo significant maturation that support improvements in cognitive control functions (Tervo-Clemmens et al., 2023). Developmental maturation of inhibitory circuitry is one proposed mechanism of suppressed spontaneous activity observed during the opening of a critical period (Hensch, 2005; Toyoizumi et al., 2013). This is in line with the known developmental decreases in spontaneous activity during resting-state (Miskovic et al., 2015; Ott et al., 2021) and the auditory steady-state response (McKeon et al., 2024a) which have been linked to reductions in behavioral response variability (Arazi et al., 2017; Arieli et al., 1996; McKeon et al., 2024a). While spontaneous neural activity is crucial for maintaining flexible neural circuitry (Toyoizumi et al., 2013), during task it may represent a neural “noise” floor, impeding response inhibition by priming individuals to make a reflexive motor response by operating close to a motor threshold (Gavenas et al., 2024; Schurger et al., 2012). Adolescents tend to have higher spontaneous activity which decreases developmentally. This may prime adolescents to make a reflexive response and therefore require greater suppression of this spontaneous activity during cognitive tasks compared to adults. This reduced condition-related suppression observed in adults (indicated by the significant age-by-condition interaction on broadband spontaneous power) may suggest that spontaneous activity is now “low enough” from a reflexive motor threshold that adults do not need to further suppress spontaneous activity during cognitive tasks to engage response inhibition, although this interpretation remains speculative. Furthermore, lower spontaneous activity during both the AS and VGS tasks are associated with reduced AS performance variability, suggesting that ongoing activity affects behavioral variability, regardless of context. The relationship between spontaneous activity and trial-to-trial latency variability was specific to frontal brain regions, suggesting that decreased spontaneous activity allows for enhanced transmission of task-evoked signals, specifically in frontal regions crucial for response inhibition (i.e. DLPFC and FEF) (Apšvalka et al., 2022; Constantinidis and Luna, 2019; Everling et al., 1998; Hwang et al., 2010; Ordaz et al., 2013). Overall, these findings provide a potential mechanism for the stabilization of inhibitory control throughout adolescence and into adulthood.

Beta oscillations have an established role in sensorimotor processing (Schmidt et al., 2019), and decreased beta power has been extensively shown to precede to movement initiation (Brown, 2007; Gehringer et al., 2019; Khanna and Carmena, 2017; Kurz et al., 2016; Little and Brown, 2014; Peles et al., 2020; Rossiter et al., 2014; Schmidt et al., 2019; Tzagarakis et al., 2010). Consistent with this literature, we found an event-related desynchronization (ERD) in the beta band during preparation of an antisaccade (AS task) and a prosaccade (VGS task). The presence of beta ERD in both conditions likely reflects the requirement of beta desynchronization in movement generation. However, the greater modulation of beta activity during AS relative to VGS may additionally reflect the increased cognitive demands needed to suppress a prepotent response and generate a voluntary antisaccade. Beyond sensorimotor processing, beta oscillations have been implicated in cognitive control functions, including working memory (Lundqvist et al., 2018, 2016; Schmidt et al., 2019) and inhibitory control (Ding et al., 2023), such that beta oscillations exert top-down control (Hwang et al., 2016) over bottom-up gamma activity (Lundqvist et al., 2018, 2016; Richter et al., 2017; Riddle et al., 2019; Trajkovic et al., 2025; Xiong et al., 2023). Adolescents showed greater modulation of beta activity during AS relative to VGS than adults, indicating that the additional demands of inhibitory preparation may exert a larger influence on beta activity earlier in development. One possibility is that immature cognitive-control circuitry requires greater task-dependent adjustment of preparatory activity to support successful inhibition, whereas mature circuitry can engage inhibitory control with less deviation from its ongoing state. Evidence has suggested that reductions in beta oscillations reduce trial-to-trial variability in neural activity (Daniel et al., 2019; Karvat et al., 2021), and, in the present study, we found a similar trend, particularly in adults. Development of beta activity may thus reflect maturational changes in prefrontal circuitry that serve to reduce levels of spontaneous activity in adulthood, allowing for improvements in both inhibitory control performance and reliability (Arieli et al., 1996; Daniel et al., 2019; Karvat et al., 2021; Schurger et al., 2012). In adolescence, beta modulation appears more closely related to task-evoked activity, whereas, in adulthood, beta modulation is more closely related to spontaneous activity. Together, these findings may reflect a developmental shift in how preparatory beta-band dynamics interact with the broader neural state.

We found that, while gamma oscillations are involved in correct and reliable AS performance, they do not appear to modulate spontaneous or evoked activity. Generally, gamma oscillations are bottom-up signals modulated by top-down beta oscillations (Bastos et al., 2020; Lundqvist et al., 2018; Richter et al., 2017; Riddle et al., 2019; Trajkovic et al., 2025; Xiong et al., 2023). Gamma oscillations are thought to support local circuit computation (Fernandez-Ruiz et al., 2023), and, in the context of movement, are theorized to be a prokinetic signal (Brücke et al., 2013; Harmony et al., 2009; Ulloa, 2022). We found that, when preparing to inhibit an eye movement, there is a large ERD (suppression) in the gamma band. Furthermore, given that gamma oscillations likely reflect bottom-up processing, we would expect these oscillations to be related to task performance (Lundqvist et al., 2018, 2016), though they may not directly contribute to the modulation of broadband spontaneous and evoked activity during task (Karvat et al., 2021). Overall, these findings suggest a role for gamma oscillations in inhibitory control engagement and its development.

In our analysis of the parietal electrodes, we found reduced spontaneous power compared to frontal electrodes during the AS task, and this difference became more prominent with age. Additionally, we found that the association between decreased spontaneous power and reduced latency variability was present only in frontal electrodes, with no significant behavioral associations with parietal spontaneous power. While we found that there were frequency-specific oscillations that contribute to AS performance, there was no modulatory effect of spontaneous or evoked power comparable to that observed for frontal high-beta activity. This is in line with the significant role of prefrontal circuitry maturation during adolescence, which supports improved cognitive performance (Larsen and Luna, 2018).

Neural activity can be characterized as periodic (oscillations) or aperiodic. Here, we examined the role of frequency-specific activity in task performance; however, we did not explicitly separate periodic and aperiodic activity. Future research should include separating out periodic and aperiodic activity during a task to determine whether the results reported here are driven more by periodic or aperiodic activity. Additionally, while our results appeared specific to frontal electrodes (as opposed to parietal electrodes), there is still the potential for effects of volume conduction in EEG analyses that make it difficult to determine whether these effects are present across the entire brain.

This study provides novel evidence that frequency-specific prefrontal dynamics contribute to the stabilization of neural processing, supporting the developmental improvements in cognitive control performance and reliability form adolescence into adulthood. Specifically, we found that beta activity, which continues to undergo maturational changes throughout adolescence, modulates ongoing activity specifically in adulthood, providing evidence that maturation of cognitive control circuitry allows beta activity to optimally exert top-down control, resulting in lower spontaneous, ongoing activity. Understanding the role of frequency-specific signaling dynamics, and their impact on intrinsic neural activity and cognitive control, may inform our understanding of psychiatric disorders that emerge in adolescence and have known alterations in oscillatory and spontaneous activity.

## Notes

https://github.com/AMB895/AS-EEG

### Competing Interest Statement

The authors have declared no competing interest.

### Funding

- National Institute of Mental Health: MH067924

