## Supplemental Material for "Developmental interactions between oscillatory and spontaneous neural activity support improvements in inhibitory control from adolescence into adulthood"

Abigail M. Beatty[1], Shane D. McKeon[1], Daniel J. Petrie[3], Finnegan J. Calabro[1,3], Ashley C. Parr[3], Will Foran[3], Beatriz Luna[1,3,4]

1. Department of Bioengineering, University of Pittsburgh, PA, USA
2. Department of Human Development and Family Science, University of Georgia, Athens, GA, USA
3. Department of Psychiatry, University of Pittsburgh, Pittsburgh, PA, USA
4. Department of Psychology, University of Pittsburgh, Pittsburgh, PA, USA

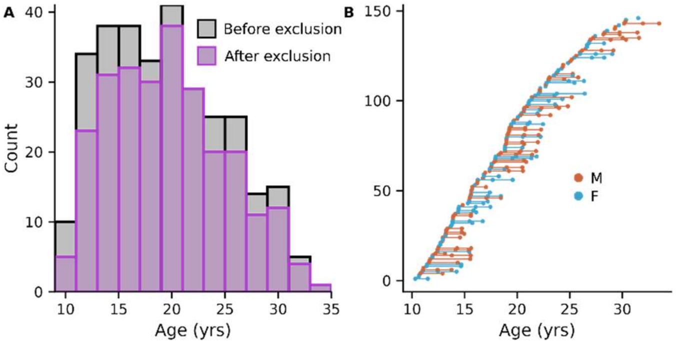


**Fig. S1: Age distribution. A.** Distribution of ages before (gray) and after (purple) exclusion. **B.** All participants included in analyses after exclusion. Each point represents a single visit and the lines connecting points (visits) for each subject. (males = orange; females = blue)

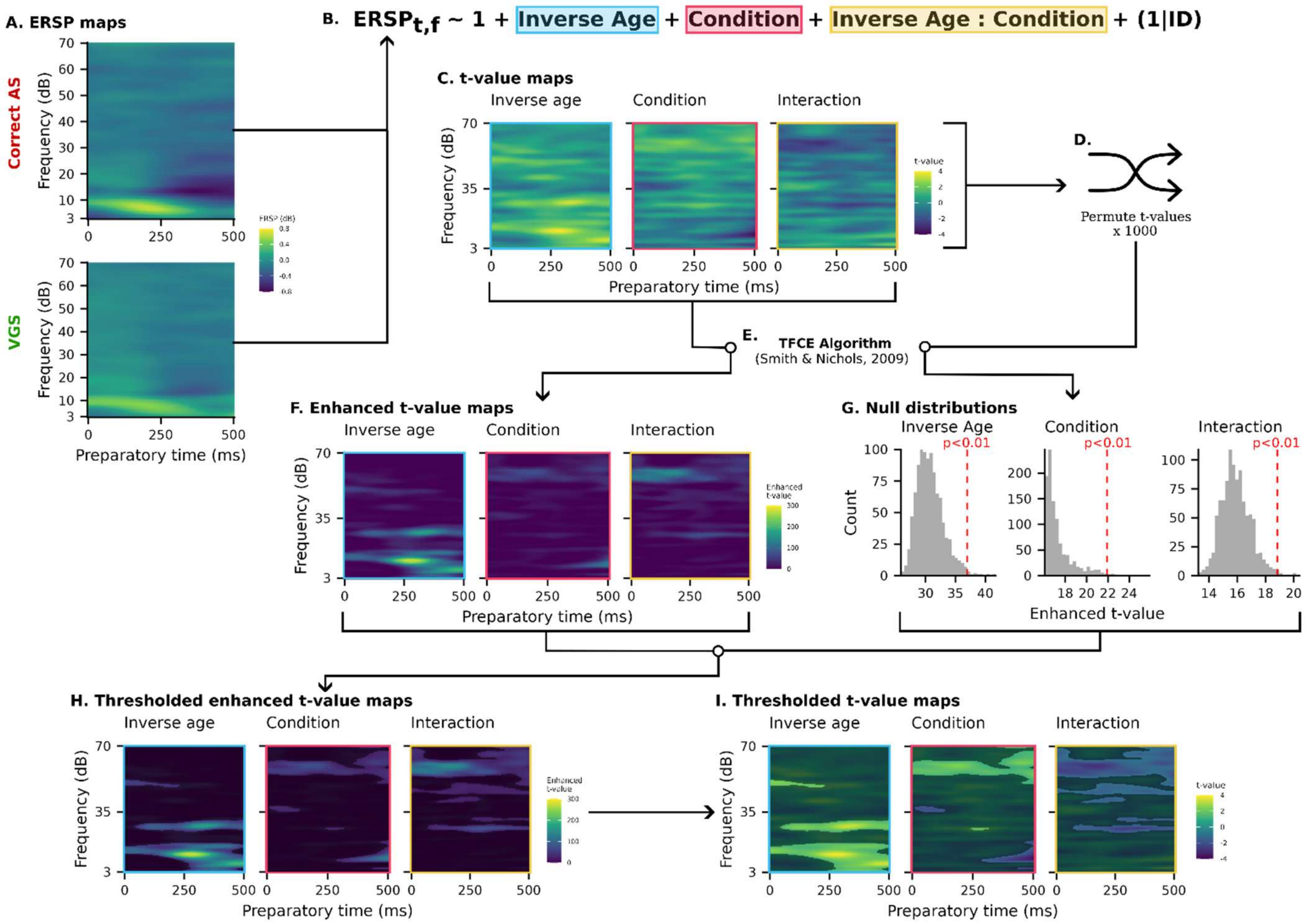


**Fig. S2: Threshold-free cluster enhancement method.** Visualization of the threshold-free cluster enhancement (TFCE) method for identifying significant time-frequency clusters. **A.** Mean event-related spectral perturbation (ERSP) maps across participants for correct AS **(top)** and VGS **(bottom)** trials. The x-axis is time during the preparatory period in milliseconds, the y-axis is frequency in Hertz, and the color bar shows the change in signal relative to the baseline period in decibels. **B.** Linear mixed effects model to estimate the effects of (inverse) age and task condition and their interaction for each time-frequency point in the ERSP maps. **C.** The t-value time-frequency maps of the main effect of age (left; blue), condition (middle; pink) and the interaction between age and condition (right; yellow) from the linear mixed effects model. The t-value maps were then input into the **E.** TFCE algorithm that **F.** enhances t-values based on strength and size of surrounding data. To identify significant clusters, the **C.** original t-value maps were **D.** permuted and input into the TFCE algorithm 1000 times. The maximum enhanced t-value from the permuted maps were used to build the **G.** null distribution and **H.** enhanced t-values greater than the $p < 0.01$ threshold of the null distribution were significant. **I.** Significant masks were then applied to the original ERSP data to extract cluster averages per subject for analysis.

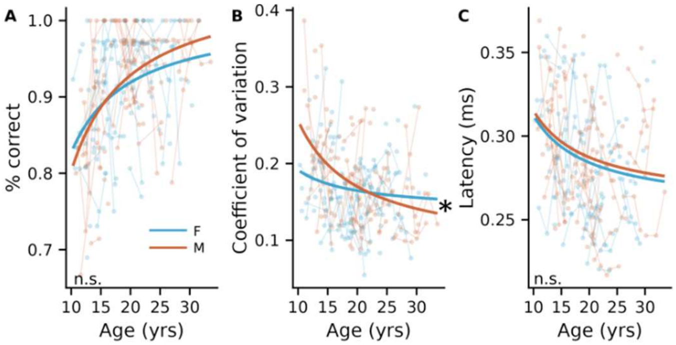


**Fig. S3: Age by sex interaction on AS behavior.** Age by sex interactions on **A.** correct response rate, **B.** latency variability, and **C.** mean latency. The asterisks represent the significance of the age by sex interaction. Only latency variability showed a significant age by sex interaction ($\beta = 1.19$, $t = 2.53$, $p = 0.04$) where males (orange) had a sharper decrease in latency variability compared to females (blue). Circles represent raw behavioral data for each subject across age, where the same subject is connected with a thin line. Lines represent the model predictions of the interaction effect of age by sex on AS behavior. (n.s. = not significant; * = $p < 0.05$)

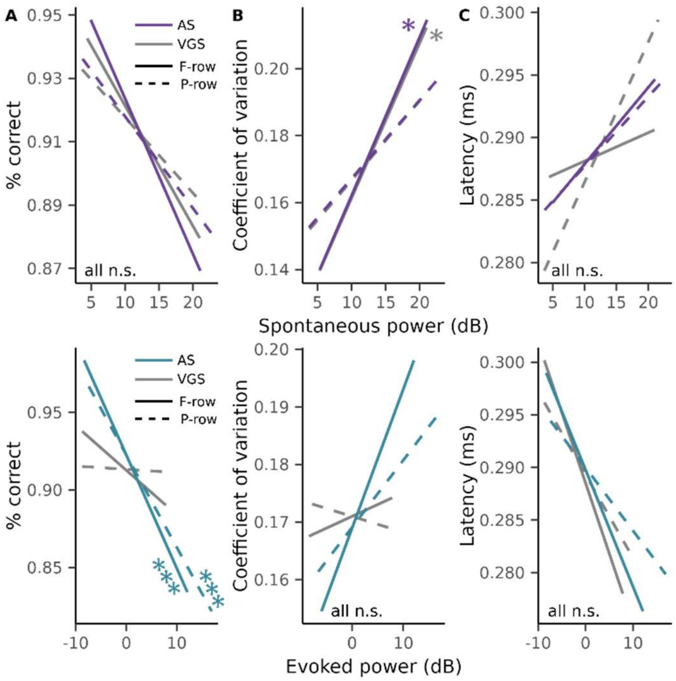


**Fig. S4: Spontaneous evoked power associations with AS behavior.** Spontaneous **(top)** and evoked **(bottom)** power associations with **A.** correct response rate, **B.** latency variability, and **C.** average latency on correct AS trials for frontal (solid lines) and parietal (dashed lines) rows and AS (spontaneous: purple; evoked: blue) and VGS (gray) tasks. Lines represent model predictions of power on AS behavior. (* = $p < 0.05$; *** = $p < 0.001$; n.s. = not significant)

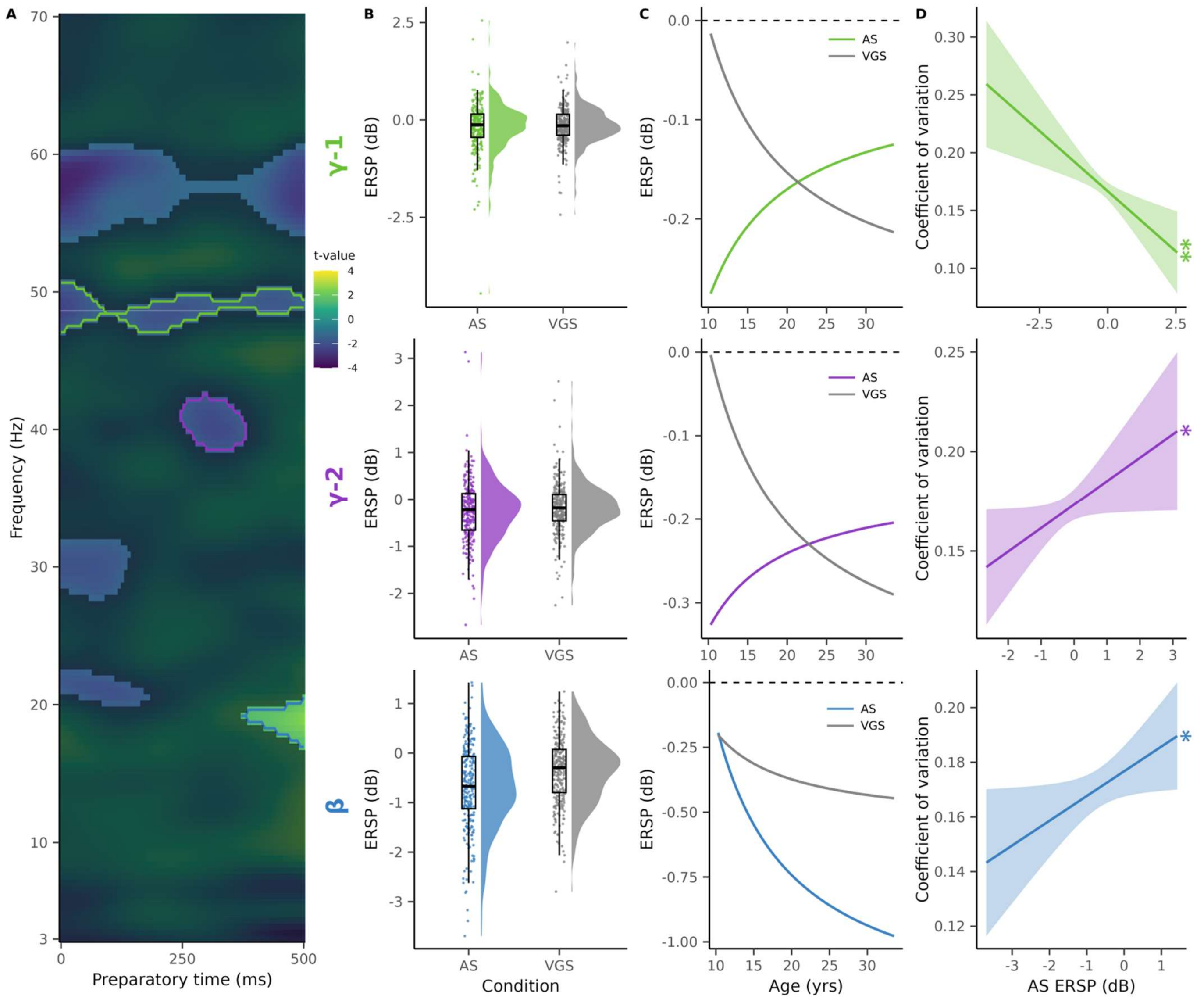


**Fig. S5: Frequency-specific oscillations associations with AS latency variability. A.** Significant parietal ERSP clusters from the interaction (age * condition) term in the linear mixed effects model from section 2.3.3.3, where the x-axis is time throughout the preparatory period in milliseconds, the y-axis is frequency and the color bar (z-axis) is the t-value. Six significant clusters where identified; however, only mean ERSP from the gamma-1 cluster (green), gamma-2 (purple), and beta (blue) were related to AS behavior. The distribution of mean ERSP from the **B. (top)** gamma-1 cluster (AS: green; VGS: gray), **B. (middle)** gamma-2 cluster (AS: purple; VGS: gray), and **B. (bottom)** beta cluster (AS: blue; VGS: gray). Due to the large variability in the clusters' mean ERSP, linear regression model predictions are shown in subsequent plots (C-D), controlling for sex (C) and age (D). Age-related trajectories of mean **C. (top)** gamma-1 cluster (AS: green; VGS: gray), **C. (middle)** gamma-2 cluster (AS: purple; VGS: gray), and **C. (bottom)** beta cluster (AS: blue; VGS: gray)**,** where the black dashed line represents no change in power from baseline, thus negative ERSP indicates a suppression in power from baseline and a positive ERSP indicates an activation in power from baseline. Associations between latency variability and **D. (top)** gamma-1, **D. (middle)** gamma-2, and **D. (bottom)** beta AS ERSP. Statistical results are reported in Supplemental Table 3. (n.s. = not significant; * = $p < 0.05$; ** = $p < 0.01$).

**Supplemental Table 1: AIC analysis with linear vs inverse age**

| **Behavior** | **Model** | **AIC** |
|---|---|---|
| **Correct response rate** | Linear age | -597.47 |
| | Inverse age | **-613.86** |
| **Latency variability** | Linear age | -698.54 |
| | Inverse age | **-707.95** |
| **Mean latency** | Linear age | -961.50 |
| | Inverse age | **-978.72** |

**Supplemental Table 1.** Summary of AIC analysis showing inverse age best models developmental improvements in inhibitory control.

**Supplemental Table 2: Age by sex interactions on spontaneous and evoked power**

| **Condition** | **Power Type** | **Row** | **Term** | **Estimate (β)** | **t-value** | ***p* (FDR corrected)** |
|---|---|---|---|---|---|---|
| AS | Spontaneous | F-row | Inverse age | 89.70 | 8.28 | < 0.001 |
| | | | Sex | -2.94 | -3.28 | 0.002 |
| | | | Inverse age*sex | 38.63 | 2.51 | 0.03 |
| | Evoked | F-row | Inverse age | 97.30 | 6.94 | < 0.001 |
| | | | Sex | -4.74 | -4.18 | < 0.001 |
| | | | Inverse age*sex | 66.09 | 3.36 | 0.008 |
| VGS | Spontaneous | F-row | Inverse age | 109.59 | 9.61 | < 0.001 |
| | | | Sex | -2.84 | -3.04 | 0.003 |

| | | | | | | |
|---|---|---|---|---|---|---|
| | | | Inverse age*sex | 36.60 | 2.26 | 0.04 |
| | Evoked | F-row | Inverse age | 115.92 | 8.41 | < 0.001 |
| | | | Sex | -2.91 | -2.60 | 0.01 |
| | | | Inverse age*sex | 39.38 | 2.03 | 0.04 |
| AS | Spontaneous | P-row | Inverse age | 106.09 | 7.82 | < 0.001 |
| | | | Sex | -3.40 | -3.06 | 0.003 |
| | | | Inverse age*sex | 39.22 | 2.05 | 0.04 |
| | Evoked | P-row | Inverse age | 131.20 | 8.08 | < 0.001 |
| | | | Sex | -4.27 | -4.94 | 0.002 |
| | | | Inverse age*sex | 66.09 | 3.36 | 0.04 |
| VGS | Spontaneous | P-row | Inverse age | 107.64 | 7.94 | < 0.001 |
| | | | Sex | -3.74 | -3.37 | 0.003 |
| | | | Inverse age*sex | 42.40 | 2.21 | 0.04 |
| | Evoked | P-row | Inverse age | 110.08 | 7.21 | < 0.001 |
| | | | Sex | -5.09 | -4.11 | < 0.001 |
| | | | Inverse age*sex | 65.31 | 3.05 | 0.01 |

**Supplemental Table 2.** Statistical results from age by sex interaction model on broadband spontaneous and evoked power for all conditions (AS and VGS) and all rows (frontal and parietal). Males tended to have lower broadband power compared to females and with a steeper decrease with age

**Supplemental Table 3: Frequency-specific oscillations associations with AS performance and reliability**

| **Electrodes** | **Cluster** | | | **Correct Response Rate** | | **Coefficient of variation** | |
|---|---|---|---|---|---|---|---|
| | | **Frequency (Hz)** | **Time (ms)** | **Estimate (β)** | ***p*** | **Estimate (β)** | ***p*** |
| **P-row** | Beta-2 | 20.1-22.3 | 0-180 | 0.009 | 0.18 | -0.007 | 0.19 |
| | Beta-3 | 27.7-31.8 | 0-140 | -0.004 | 0.62 | -0.006 | 0.24 |

| | | | | | | |
|---|---|---|---|---|---|---|
| Gamma-1 | 47.1-50.7 | 0-500 | 0.02 | 0.06 | **-0.02** | **0.001** |
| Gamma-3 | 53.8-60.6 | 0-500 | -0.0006 | 0.95 | 0.009 | 0.20 |
| Gamma-2 | 38.5-42.6 | 247-380 | 0.003 | 0.73 | **0.01** | **0.05** |
| Beta-1 | 16.9-20.5 | 373-500 | 0.001 | 0.84 | **0.009** | **0.04** |

**Supplemental Table 3.** Summary of AS ERSP clusters and their associations with behavior. F-row corresponds to electrodes associated with parietal brain regions.
